# Long-term 2D monoculture of primary mouse LSEC preserves scavenging capacity and enables siRNA knockdown of Mrc1

**DOI:** 10.64898/2026.05.04.722602

**Authors:** K. Szafranska, B. Abujayyab, E. Struck, D. Hovland, C. Holte, G. Dumitriu, K Sørensen, P. McCourt

**Affiliations:** Vascular Biology Research Group, Dept. Medical Biology, University of Tromsø UiT The Arctic University of Norway, Tromsø, Norway; Translational Vascular Research Group, Dept. of Clinical Medicine, University of Tromsø UiT The Arctic University of Norway, Tromsø, Norway

**Keywords:** liver sinusoidal endothelial cells (LSEC), long-term primary cell culture, endocytic capacity, in vitro siRNA knockdown, Mrc1 knockdown, hepatic sinusoids

## Abstract

Liver sinusoidal endothelial cells (LSEC) rapidly dedifferentiate in 2D-monoculture, losing their high endocytic activity and characteristic morphology, limiting their use in mechanistic studies. We established and validated culture conditions that preserve LSEC endocytic capacity for at least 10 days, enabling efficient in vitro siRNA-mediated gene silencing. Mouse LSEC were cultured in 5% oxygen, growth media partially exchanged daily and assessed for cell viability, endocytic capacity, morphology and ultrastructure. Despite typical culture-induced defenestration, the cells showed high viability and efficient endocytosis via scavenger-receptors. This allowed for siRNA-mediated mannose receptor knockdown exemplified by 96% and 76% reduction in Mrc1 mRNA and protein expression at 72h (validated by qPCR and Western blot), with functional assays confirming decreased mannose-receptor-mediated endocytosis. Extended maintenance of LSEC viability and functions, previously restricted to complex co-culture systems, provide a practical platform for investigating LSEC-specific molecular mechanisms and hepatic sinusoid physiology.

## 1. Introduction

The walls of hepatic sinusoids are lined with highly specialised endothelial cells, essential for normal liver functioning. The liver sinusoidal endothelial cells (LSEC) show a unique morphology with numerous fenestrations – transcellular patent pores with diameters of 50-350 nm [1], [2], which provide passive, bidirectional and size-dependent exchange of macromolecules between circulating blood and hepatocytes. In addition, LSEC have the highest scavenging capacity among the body’s endothelia and express a set of endocytosis receptors, which are rare for other vascular beds, such as stabilin-1, stabilin-2, the mannose receptor (MRC1, CD206) and Fc-gamma receptor IIb2 [3]. These scavenging receptors operate via clathrin-mediated endocytosis and allows for the active removal of waste macromolecules [4], [5], such as connective tissue molecules from tissue turnover [6], [7], oxidized and glycated plasma proteins [8], [9], [10], and viruses and virus-like particles [11], [12], [13], therefore contributing to the maintenance of blood and liver homeostasis [14], [15], [16].

These unique morphological and functional features make LSEC highly specialised and differentiated endothelial cells. *In vivo*, LSEC are heavily influenced by paracrine and autocrine signalling from hepatocytes and other liver cells, as well as mechanical cues from the physiologically soft extracellular matrix of the liver sinusoid [17]. These conditions are extremely difficult to replicate outside of the native liver environment which makes the long-term *in vitro* culture of primary LSEC particularly challenging. Primary LSEC typically lose their fenestrations, high endocytic activity as well as other cell-specific markers within a few days in culture, resulting in phenotypic changes that make them unsuitable for extended studies [18][19][20]. This rapid dedifferentiation poses a critical concern for reproducibility [21], as studies utilising LSEC beyond the dedifferentiation window without rigorous ultrastructural and functional validation may inadvertently report data from generic capillary-type endothelium rather than true sinusoidal endothelial cells. Attempts to establish an LSEC cell line encounter similar challenge resulting in several LSEC-like cell lines preserving only selected specific functions [22].

The sinusoidal microenvironment can be partially recreated *in vitro* by 2D and 3D co-culture systems that combine several types of liver cells, such as hepatocytes, hepatic stellate cells and liver macrophages [23], [24]. These systems, in many aspects, outperform monocultures and are particularly valuable for *in vitro* hepatotoxicity assessment where interactions between different cell types can be crucial to determine toxicity mechanisms. Although a few studies reported successful maintenance of some of the main LSEC functions for several days or even weeks [25], [26], [27], the cellular complexity of co-cultures introduces practical limitations for mechanistic studies. Specifically, the presence of multiple cell types prevents or complicates the use of downstream analyses that require pure, homogeneous cell populations, such as RNA-based gene silencing. For example, *in vitro* siRNA mediated knockdown can be crucial for revealing the still unknown molecular architecture of LSEC fenestrations or identifying novel regulators of the scavenging system. Both applications would require prolonged survival and transfection efficiency of a pure LSEC population, ideally maintained for several days post-transfection to allow protein turnover and phenotypic responses.

Therefore, in this study we optimised and performed a thorough characterization and validation of cell-type specific functions of mouse LSEC kept under optimized culture conditions in a 2D monoculture system for up to 10 days with the dual goal of (1) characterising the temporal dynamics of the LSEC phenotype and (2) establishing a platform for *in vitro* siRNA-mediated gene silencing. We show that the culture conditions secured highly reproducible preservation of the LSEC scavenging capacity, during the whole experimental period, allowing a successful siRNA knockdown of Mrc1, enabling functional validation of receptor-dependent endocytosis and providing a foundation for future studies of LSEC-specific molecular regulators using siRNA.

## 2. Materials and methods

### 2.1. Animal ethics

The experiments followed protocols approved by the competent institutional authority at UiT-The Arctic University of Norway, licenced by the National Animal Research Authority at the Norwegian Food Safety Authority (Approval IDs: UiT 09/22, 12/23, 13/24). The animal experiments (i.e., liver perfusions for cell isolation) were performed postmortem, and the mice were euthanized by cervical dislocation before the start of the procedure. All experiments were performed in accordance with relevant approved guidelines and regulations and reported following ARRIVE guidelines. Each biological replicate consisted of cells from one animal, except in Western blot experiments (section 2.9.) where each biological replicate contains pooled cells from two animals to achieve the required protein concentration per sample. All experiments were performed in a minimum of 3 biological replicates with the total use of 45 mice.

### 2.2. Primary LSEC isolation and culture

LSEC were isolated as described in [28]. Briefly, liver cells were isolated from adult male C57BL/6JRj mice (Janvier Labs, France, 3-4 months old) by liver perfusion via portal vein with Liberase TM (Roche, Germany). LSEC were separated using low-force differential centrifugation followed by a magnetic activated cell sorting system (MACS, Miltenyi Biotec, Germany) with mouse anti-CD146 beads. Culture purity was assessed using scanning electron microscopy (SEM) showing >95% of the cells displaying fenestrated morphology (LSEC hallmark).

After isolation, LSEC were seeded in a close to monolayer confluency (300 000 cells/cm^2^) on fibronectin coated (15 min coating time in room temperature (RT) with 0.2 mg/mL of human plasma fibronectin) tissue culture plastic 12/24/48-well plates (Sarstedt) in Endothelial Growth Media (EGM, cat# 211-500, Cell Applications, USA). According to manufacturer’s information EGM contains: 2% fetal bovine serum, 1 g/L glucose, penicillin, streptomycin, and Amphotericin B, hydrocortisone, heparin and growth factors excluding VEGF. Cells were cultured in 37°C, 5% CO_2_, 5% O_2_ with half of the media volume exchanged daily. “Day1” is the day of the LSEC isolation, and any experiments were started about 4h after cell seeding. The cells were evaluated on days 1,3,5,8,11 as indicated in each result section.

### 2.3. Scanning Electron Microscopy

LSEC seeded on 24-well plates were fixed on days 1,3,5 in a mix of 4% formaldehyde (FA) and 1% glutaraldehyde (GA) (in PHEM buffer) for 10 min in RT. Samples were then stored in fixative at 4°C until post-processing. After replacing fixative with PHEM buffer, bottoms of the plastics were detached using a mechanical drill and samples were then post fixed for 1h in 1% tannic acid in PHEM buffer and 30 min in 1% osmium tetroxide in double distilled H_2_O, then by dehydrated in an ethanol gradient (5 min of 30/60/90% and 4x 5 min of 100% ethanol) followed by chemical drying with hexamethyldisilazane (HMDS, cat# 999973) for 2x 5 min. Samples were mounted on aluminium stubs, stored in a desiccator at RT and sputter coated with 10 nm of gold/palladium alloy directly before imaging.

Overview SEM images were taken using Zeiss Gemini300 at magnification 300x, 2keV, 20nm pixel size. Images were taken from 4-5 separate areas from each sample and total of 500-1000 cells per sample were assessed by semi quantitative analysis (described in detail previously in Supplementary Information of [29]). Briefly, all cells were manually assigned based on their morphology: highly/normally/low fenestrated and defenestrated.

### 2.4. Light microscopy

Phase contrast images of LSEC seeded in 48-well plates were taken from same samples/wells on days 1, 3, 5, 8 and 11 with a 40x objective using an EVOS (ThermoFisher Scientific) inverted microscope.

### 2.5. Fluorescence microscopy

LSEC were cultured in 48-well plates until fixation in 4% FA in PBS for 10 min at RT. Samples were stored at 4°C in 0.1% FA in PBS until staining. First, samples were permeabilized by incubation for 2 min in 0.05% Triton X-100, then washed 3× with PBST (PBS with 0.5% Tween20) and incubated in the dark overnight at RT with 0.01 mg/mL anti-tubulin-AF647 monoclonal antibody (sc-23948, Santa Cruz Biotechnology, USA). Then, 50 µg/mL phalloidin-AF555 (A34055, Invitrogen, ThermoFisher, USA) in PBST was added for 30 min and 2 mg/mL 4’,6-diamidino-2-phenylindole (DAPI) (Sigma, USA) for 15 min. Finally, samples were washed with PBST 3×10 min followed by 2×30 min. Fluorescent images were taken using an EVOS (ThermoFisher Scientific) inverted microscope.

### 2.6. Viability

#### 2.6.1. Cell number

LSEC were seeded in 48-well plates and stained with either DAPI or Hoechst 33258 to visualize the cell nuclei. Images were taken using EVOS (10x objective), cell nuclei were counted using a threshold-based Fiji [30] macro and the total number of cells per well was calculated. From each sample from 10 bioreplicates (each with at least 2 technical replicates), 6-8 images across the whole well were acquired and analysed.

#### 2.6.2. Resazurin assay

LSEC were seeded in 48-well plates and a commercial resazurin/resorufin assay (Abcam, Cat# ab129732) was used as an indicator of the functional viability. On selected timepoints (day 1,3,5,8 and 11) cells were supplied with 1:10 resazurin reagent to the culture media and incubated for 2.5 h. Then, cell media (50 µL) was collected and the fluorescence intensity measured with a CLARIO star microplate reader (BMG Labtech, excitation/emission of 545-20/600-40 nm).

#### 2.6.3. LDH assay

LSECs were seeded on 48-well plates and a luminescence lactase dehydrogenase (LDH) detection kit (LDH-Glo, Promega cat#J2380) was used following the manufacturer’s instructions to assess cell viability. On selected timepoints (day 1, 3, 5, 8 and 11) cells were lysed for 5 min in 1% SDS solution and 50 µL samples were collected into 450 µL of freezing buffer (details in the manufacturer’s protocol) and stored at −20°C until measurement.

### 2.7. Endocytosis assays

#### 2.7.1. Endocytic capacity study

On the day of the experiment samples were supplied with fresh EGM media containing 1% human serum albumin (Alburex 20, CSL Behring, UK), approximately 50 ng/mL of ^125^I-FSA (formaldehyde treated serum albumin [31]) or ^125^I-RibB (ribonuclease b, Sigma cat# R7884) combined with different concentrations of unlabelled FSA (0-120 µg/mL) or RibB (0-200 µg/mL) and incubated for 2h. The supernatant was transferred into tubes with the same volume of 20% trichloroacetic acid and centrifuged to separate the precipitated proteins from soluble fraction. Cells were then incubated for 20 min with 1:1000 Hoechst 33258 in EGM to stain the nuclei and 6-8 images were taken for each well (consecutive images covering the centreline of the whole well) to calculate total number of cells per well. Immediately thereafter, the cells were lysed with 1% sodium dodecyl sulfate (SDS) and transferred to tubes for radiation measurement (gamma counter Cobra II, Packard).

The amount of intracellular degradation products released after endocytosis was determined by measuring acid-soluble radioactivity in the supernatant and subtracting the acid-soluble radioactivity (representing free iodine) in the supernatant of cell-free control wells. Cell-associated ligand was quantified by measuring SDS soluble radioactivity in the remaining cells minus the radioactivity of nonspecific surface binding in cell-free control wells. All experiments were performed in 6 biological replicates, each in 3 technical replicates per experiment. All data was then normalized to the number of cells per well.

#### 2.7.2. Endocytosis of fluorescent ligands

LSEC cultures in 48-well plates were treated for 1h with one of the following scavenging ligands (in EGM media), FSA (10 μg/mL), RibB (40 μg/mL), aggregated gamma globulin (AGG, 40 μg/mL) conjugated with either AlexaFluor-488 or -647 and then rinsed with pre-warmed media to remove excess ligands before fixation with 4% FA in PBS for 15 min at RT. Fluorescent images were taken using an EVOS inverted microscope.

### 2.8. Gene knockdown/RT qPCR

Gene knockdown was performed using siRNA transfection targeting MRC1 (cat# 4390771, ThermoFisher). Transfections were carried out in OptiMEM (ThermoFisher) using Lipofectamine RNAiMAX (ThermoFisher) following the manufacturer’s protocol. siRNA (30 pmol, corresponding to 3 µL) was mixed with 9 µL Lipofectamine RNAiMAX in a total volume of 1.3 mLmedium and incubated for 5 min to allow complex formation. Cells were maintained under identical conditions, and confluency and cell morphology were monitored to ensure comparable physiological states between knockdown and control groups.

Knockdown efficiency was assessed by RT qPCR (TaqMan chemistry) using the ΔΔCt method. 18S rRNA served as internal reference and was quantified with a VIC labelled primer–probe set (cat# 4310893E, ThermoFisher). Target gene assay for MRC1 (Mm01329359_m1, ThermoFisher) were conjugated to FAM probes to be analysed simultaneously in the same well. Gene expression was measured 24h and 72h after transfection, corresponding to study days 3 and 5.

Differences between untreated and scrambled controls were evaluated first. When the scrambled control did not differ from untreated cells (fold change < 10 percent), statistical testing focused only on the comparison between knockdown and scrambled control. All conditions were measured across the same 4 biological replicates. Group differences were analysed using a linear mixed effects model with condition as fixed effect and biological replicate as random intercept. For visualization and calculation of knockdown efficiency, ΔCt values were converted to relative expression using the 2^(-ΔΔCt) method, and knockdown efficiency was calculated as one minus the ratio of mean knockdown to mean scrambled expression.

### 2.9. Western blotting

LSEC cultures in fibronectin-coated 12-well plates (~3 million LSEC per well) were transfected with siRNA as described in section 2.7. At 24h and 72h post transfection, LSEC were collected in RIPA lysis buffer (ThermoFisher, cat# 89900) containing protease inhibitor cocktail (Roche), vanadate, pepstatin A and NEM. Total protein concentration was measured with Pierce™ Detergent Compatible Bradford Assay Kit (ThermoFisher, cat# 1863028,). Sonicated samples were reduced, heated at 70°C for 10 min and loaded on SDS-Page with NuPage 3-8% Tris-acetate gels (Invitrogen) according to the manufacturer’s protocol with the recommended protein ladders (HiMark Pre-stained protein standard (cat# LC5699, Invitrogen) together with Himark Unstained protein standard (cat# LC5688) and MagicMark Western protein standard (cat# LC5603, Invitrogen) for high range molecular weight. Immunoblotting was performed onto a 0.45 µm PVDF transfer membrane (ThermoFisher, cat# 88518). The membrane were blocked with EveryBlot Blocking Buffer (Bio-RAD, cat# 120110020) for 5-10 min at RT, labelled with 2 µg/mL Goat Anti Human MMR/CD206 polyclonal antibody (R&D Systems, cat#, validated for mice in [32]) and 0.5µg/mL GADPH rabbit polyclonal antibody (Invitrogen, cat# PA1-16777) for 1h at RT, followed by three washes with TBS-T, then incubated with 0.1 ug/mL with Donkey anti-Goat IgG H&L (HRP) (Invitrogen, cat# A16005) and 0.015 ug/mL Goat Anti Rabbit IgG H&L(HRP) (Abcam, cat# ab205718, 1/10000 dilution). The proteins of interest were visualized with Super SignalWest Pico plus chemiluminescent substrate (Thermo Scientific, product# 34580) and ImageQuant LAS4000. ImageJ was used for the relative quantification of the protein bands.

### 2.10. Further statistical analyses

For RT qPCR and Western blot data, all measurements were converted to relative expression after normalisation to their respective reference signals. RT qPCR data were analysed using 2^(-ΔCt) values, and Western blot signals were normalised to the loading control. Group differences were evaluated with linear mixed effects models that included condition as fixed effect and biological replicate or blot as random intercept. When untreated and scrambled controls did not differ, statistical testing focused on the comparison between the knockdown and the scrambled control. Normality of models was assessed using Shapiro Wilk tests.

For viability assays, normality at each time point was assessed using Shapiro Wilk tests. Time dependent changes were evaluated with linear regression to estimate overall trends, followed by pairwise contrasts with the earliest measured time point and between all individual time points. Segmented regression was used to identify potential breakpoints that indicated shifts in the temporal response pattern. Relevant significance levels are indicated on in the figures and/or in the text. All statistical analyses were performed in R (Version 5.4.1).

## 3. Results

Primary mouse LSEC were isolated and cultured for up to 10 days. The preservation of LSEC endocytic capacity allowed for the demonstration of RNA-based *in vitro* knockdown of mannose receptor (MRC1). Additionally, LSEC scavenging functions, viability, morphology and ultrastructure were assessed in long-term culture.

### 3.1. Endocytic capacity is preserved in long-term culture of primary LSEC

The high-throughput scavenging system is one of the hallmarks of LSEC. It combines rapid endocytosis, a distinct set of high affinity receptors with extremely fast turnover rate as well as high content of lysosomal enzymes allowing for efficient catabolism of endocytosed material [3]. This endocytic function of primary LSEC cultured for up to 11 days was studied using radiolabelled and fluorescently labelled ligands – FSA, RibB, AGG, ligands of stabilin-1/-2, mannose receptor and FcγIIb2, respectively (**Figure 1**).

**Figure 1.**
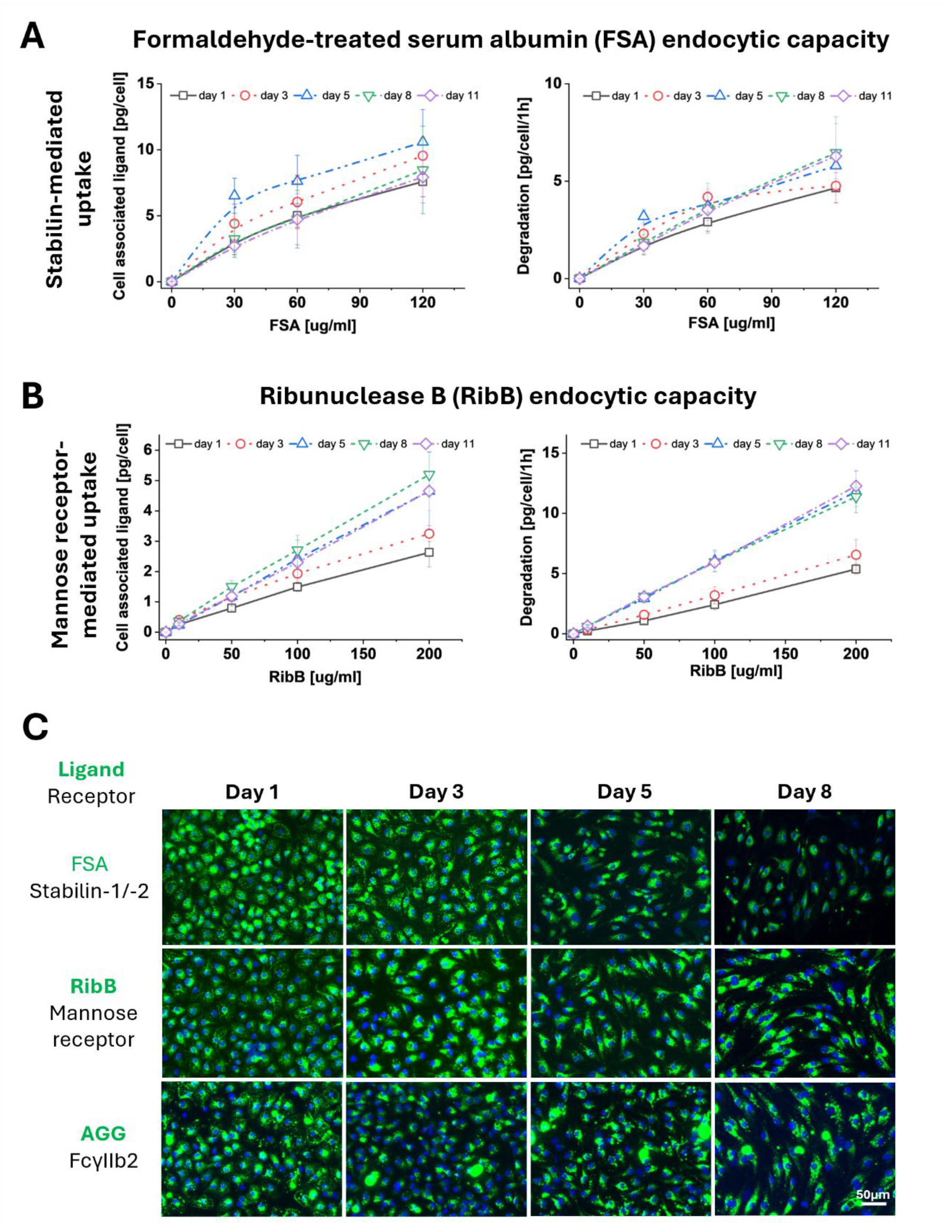
Preservation of scavenging function in mouse LSEC in vitro. Endocytic capacity in LSEC cultured for up to 10 days was assessed using radiolabelled **(A)** formaldehyde-treated serum albumin (FSA) and **(B)**, ribonuclease B (RibB), ligands of (primarily) stabilin-1/-2 and mannose receptor, respectively. Cell cultures were incubated with 0-200 μg/mL of nonlabelled ligand and ~50ng/mL of ^125^I-radiolabelled ligand (tracer) and radioactivity of cell associated and degraded fractions were measured and analysed normalizing to the number of cells in each well (n=6), mean±SD. Statistical analysis are provided in Supplementary Information Figure S1. **(C)** Endocytic activity of the whole population of LSEC was studied using fluorescently labelled ligands (green), cell nuclei (blue, DAPI). Cells were incubated for 2h with 10 μg/mL of FSA-AF488, 40 μg/mL RibB-AF647 or 40 μg/mL AGG-AF647 (aggregated gamma globulin, ligand of FcγRIIb2).

The quantitative assessment of the LSEC uptake of ^125^I-FSA and ^125^I-RibB (AUC analysis, Supplementary Information Figure S1) showed that the endocytic capacity is maintained during the whole experimental period of 10 days. The cell-bound fraction of FSA (**Figure 1A**) steadily increased, at day 5 reaching a peak of 150% relative to day 1 (p = 0.014) and later from day 8 decreasing to the level observed on day 1. The degradation efficiency for FSA did not significantly change during the experiment. The highest level was detected at day 5 although this increase did not reach statistical significance versus day 1 (p = 0.099).

Lysosomal enzymes contain terminal mannose groups and LSEC partly depend on mannose receptor-mediated recruitment of these enzymes for high degradation capacity [33]. The LSEC endocytic capacity of ^125^I-RibB (**Figure 1B**) initially increases, at day 8 reaching peak of 189% relative to day 1 (p < 0.001) for cell associated fraction of RibB. The RibB degradation capacity increased initially, at day 5 reaching 237% of level observed at day 1 (p < 0.001) and then stabilizing at the same level at day 8 (p = 0.91) and day 11 (p = 0.92).

To verify whether the LSEC scavenging function is preserved among the whole cell population or represent a minor fraction of “super-efficient” cells, we incubated LSEC cultures with fluorescently tagged ligands (FSA-AF488, RibB-AF647, AGG-AF647) for 2h at days 1,3,5 and 8 (**Figure 1C**). Similar uptake results were observed for all ligands with nearly all cells presenting fluorescent signal from day 1 to day 8, however, the ligand distribution within the cells changed with time. On day 1, the ligands were uniformly distributed in the whole cell, while in the following days, the signal was mainly localised in the perinuclear area.

### 3.2. *In vitro* siRNA knockdown of the mannose receptor

The preservation of scavenging function in long-term cultured LSEC allowed us to demonstrate the *in vitro* siRNA-based knockdown of mannose receptor (MRC1). The cells were transfected 18h after isolation (day 2) and knockdown efficiency was evaluated by measuring Mrc1 RNA and protein expression 24h and 72h post transfection, respectively (on day 3 and 5), as well as by assessment of endocytic capacity of RibB 72h post transfection (on day 5) (**Figure 2**). No significant differences in cell confluency or viability were observed between knockdown and control groups of the same day.

**Figure 2.**
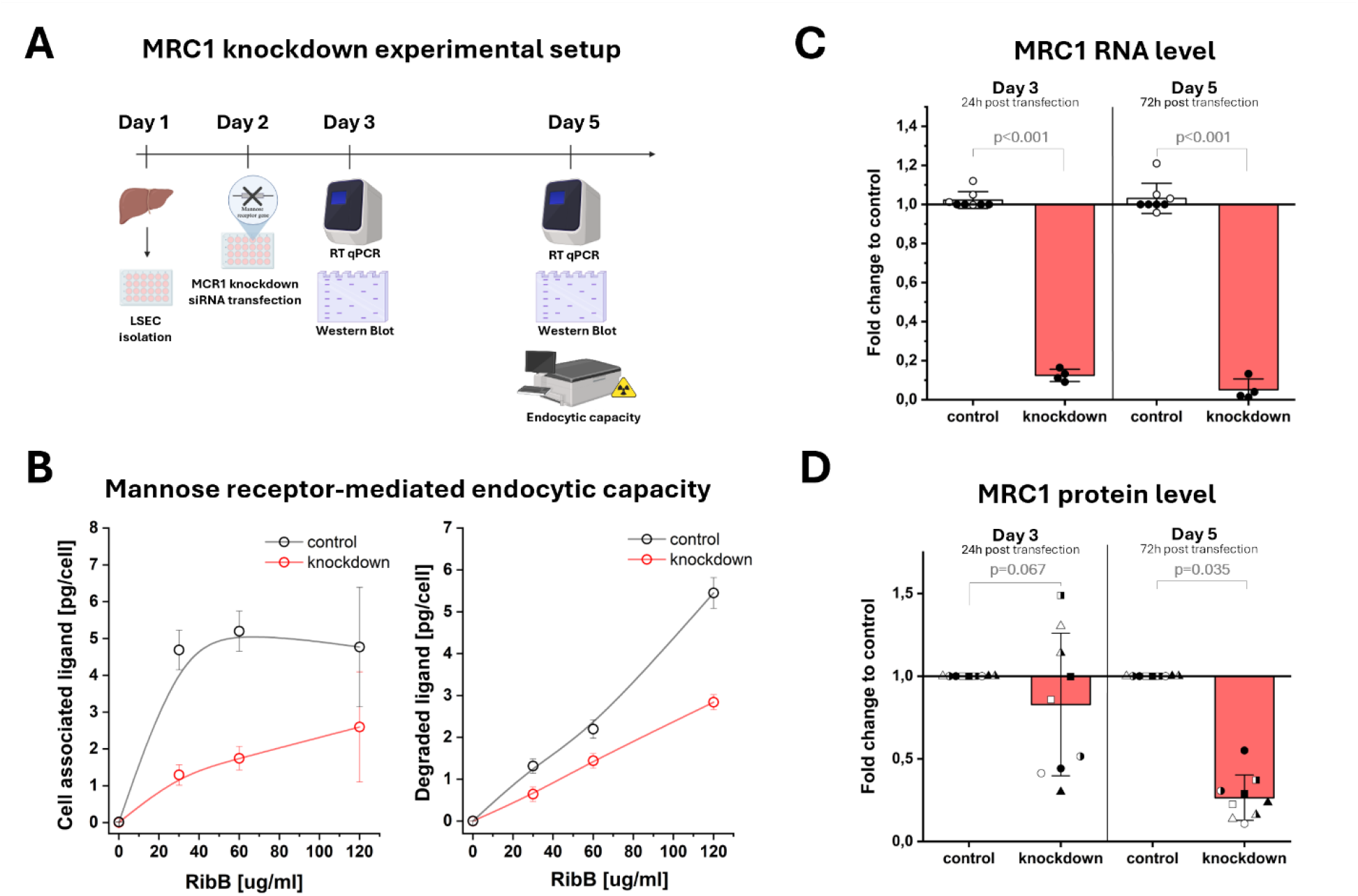
In vitro siRNA-based knockdown of MRC1 in mouse LSEC. **(A)** Schematic representation of the experimental timeline. **(B)** Mannose receptor-mediated endocytic capacity was evaluated 72 h post transfection (on day 5 of culture) using radiolabelled ribonuclease b (^125^I-RibB). Cell associated and degraded fractions were normalised to the number of cells in each well (n = 3). **(C/D)** Cells were transfected within 24h after isolation, and MRC1 protein and RNA levels were examined at 24h and 72h post transfection. **(C)** RT qPCR data were analysed using 2^(-ΔCt) values and normalised to the appropriate control. Differences between conditions were tested with linear mixed effects models. White dots represent additional untreated controls (n = 4). **(D)** Knockdown efficiency was additionally assessed by Western blot (raw data in supplementary information Figure S2), protein signals were normalised to the appropriate control and analysed with linear mixed effects models (n = 9, indicated by datapoint shape/colour). All data is shown as mean ± SD.

Mrc1 expression at RNA level was effectively reduced to 11% after 24 h and to 4% after 72 h (both p < 0.001, **Figure 2C**). Protein levels were reduced to 71% after 24 h (p = 0.067) and 24% after 72 h post transfection (p = 0.035, **Figure 2D**). The rather weak effect observed 24 h after transfection is consistent with previously described Mrc1 protein turnover, since receptor molecules synthesised just before transfection are still likely present at this early time point. Prior work reports a half-life of more than 30 h for the mannose receptor [34].

Moreover, the functional endocytic capacity assay with radiolabelled RibB as ligand showed nearly 70% reduction in cell-associated ligand 72h after knockdown which matches the Mrc1 protein expression (**Figure 2B**). Furthermore, only about 30% reduction of degradation capacity was observed, even though LSEC partly depends on mannose receptor-mediated uptake of lysosomal enzymes to keep up their high endocytic capacity in vivo [33].

Together, the consistent reduction in RNA expression, protein abundance, and receptor-mediated endocytosis confirms that the knockdown is both efficient and functionally meaningful and that the established long-term culture of mouse LSEC, together with the siRNA protocol, can be successfully applied *in vitro* to silence genes of interest.

### 3.3. Viability parameters of long-term culture of LSEC

To further characterise the LSEC in long-term culture, we assessed the cell viability parameters (**Figure 3**). The cell number steadily decreased and stabilised after day 8 at about 30% relative to day 1 (**Figure 3A**). The confluency of the cell culture remained at nearly 100% monolayer characteristic for endothelial cells (as shown in microscopy images in **Figure 4** and **Figure 5**), which suggests that the size of the individual cells increased. This observation correlated with the gradual 2.5-fold increase in the intracellular LDH level, which later stabilises from day 5 (**Figure 3C**). The functional viability assessment with resazurin/resorufin assay allowed for estimation of the reducing potential of the cell. On day 3 and day 5, we observed an increase in resazurin reduction to 175% of the initial day 1 level, which later gradually decreased, reaching 70% on day 11 (**Figure 3B**). The elevated reducing potential of the cell may reflect the response to oxidative stress stimuli in the culture, resulting from cell death, as shown in decreasing cell number. We further investigated the culture-induced changes in LSEC morphology (**Figure 4A**), cytoskeleton (**Figure 4B**) and ultrastructure (**Figure 5**).

**Figure 3.**
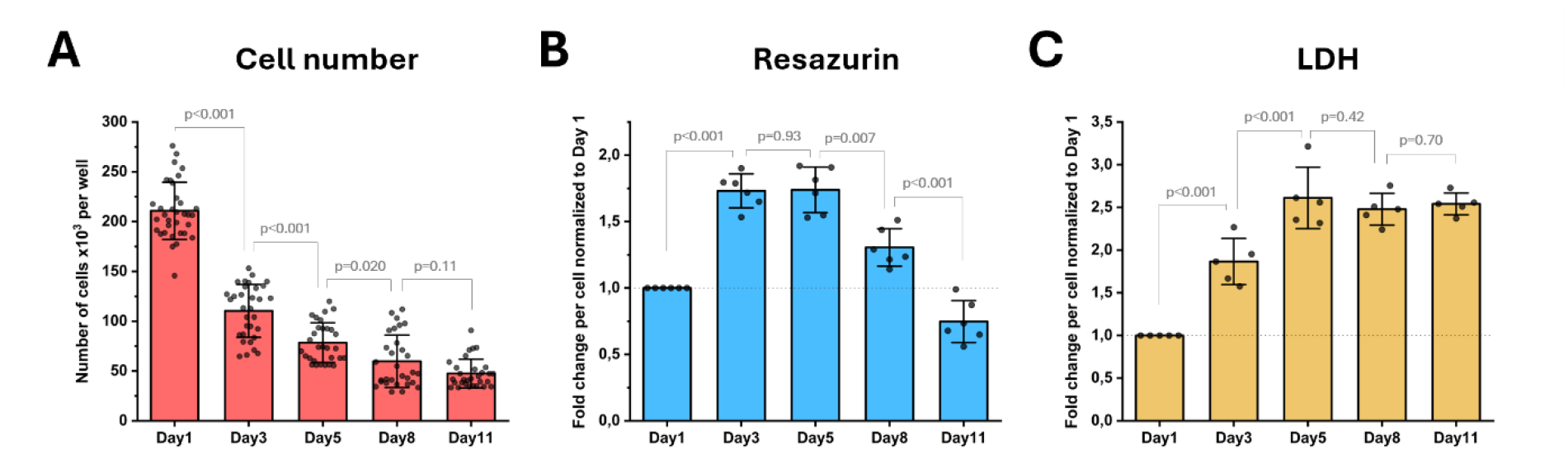
Viability parameters of primary mouse LSEC during 10 days in culture. **(A)** Cell number was calculated based on the nuclei staining with Hoechst 33258 (n=12). **(B)** Functional viability and reduction capability of the cell was studied using resazurin assay, the results were normalised to the cell number and further normalised to day 1 (n=6). **(C)** Intracellular LDH levels were measured to assess the structural integrity and cellular volume (n=5), the results were normalised to the cell number and further normalised to day 1. Each point represents a single well **(A)** or a single bioreplicate **(B**,**C)**. Data are shown as mean ± SD. Significance was evaluated using linear regression for temporal trends and pairwise contrasts between time points.

**Figure 4.**
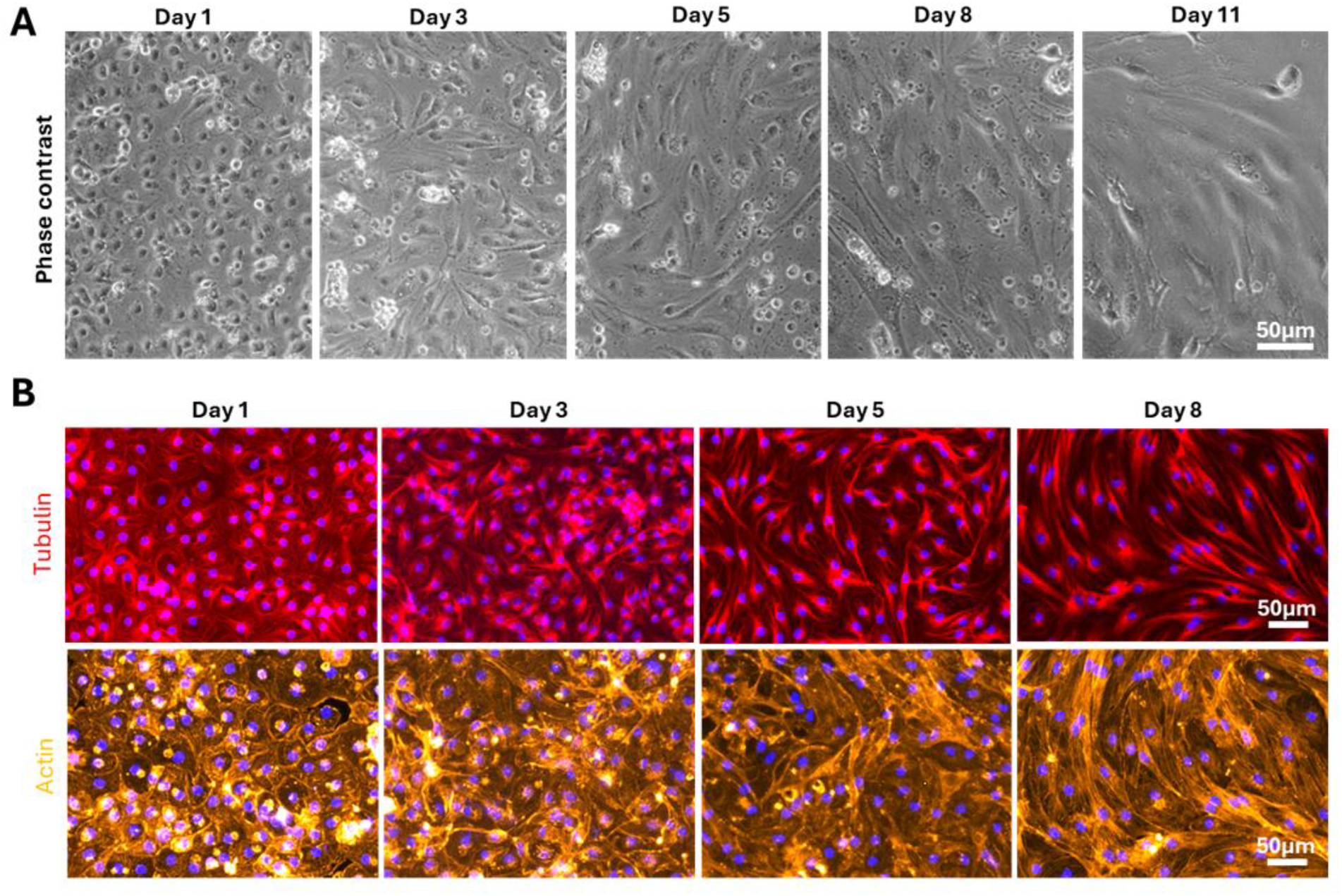
Changes in the morphology of primary mouse LSEC in culture. The general morphology of the cultures was studied using phase contrast microscopy **(A)** while actin and tubulin cytoskeleton **(B)** was stained using phalloidin-AF555 and antibody anti-α-tubulin-AF647, respectively. The confluency of the culture is maintained with increasing cell area, despite the decrease in the cell number. The shape of the cells also changes to more elongated towards day 8, with a dense actin and tubulin cytoskeleton. The size of the cell nuclei remains unchanged in culture.

**Figure 5.**
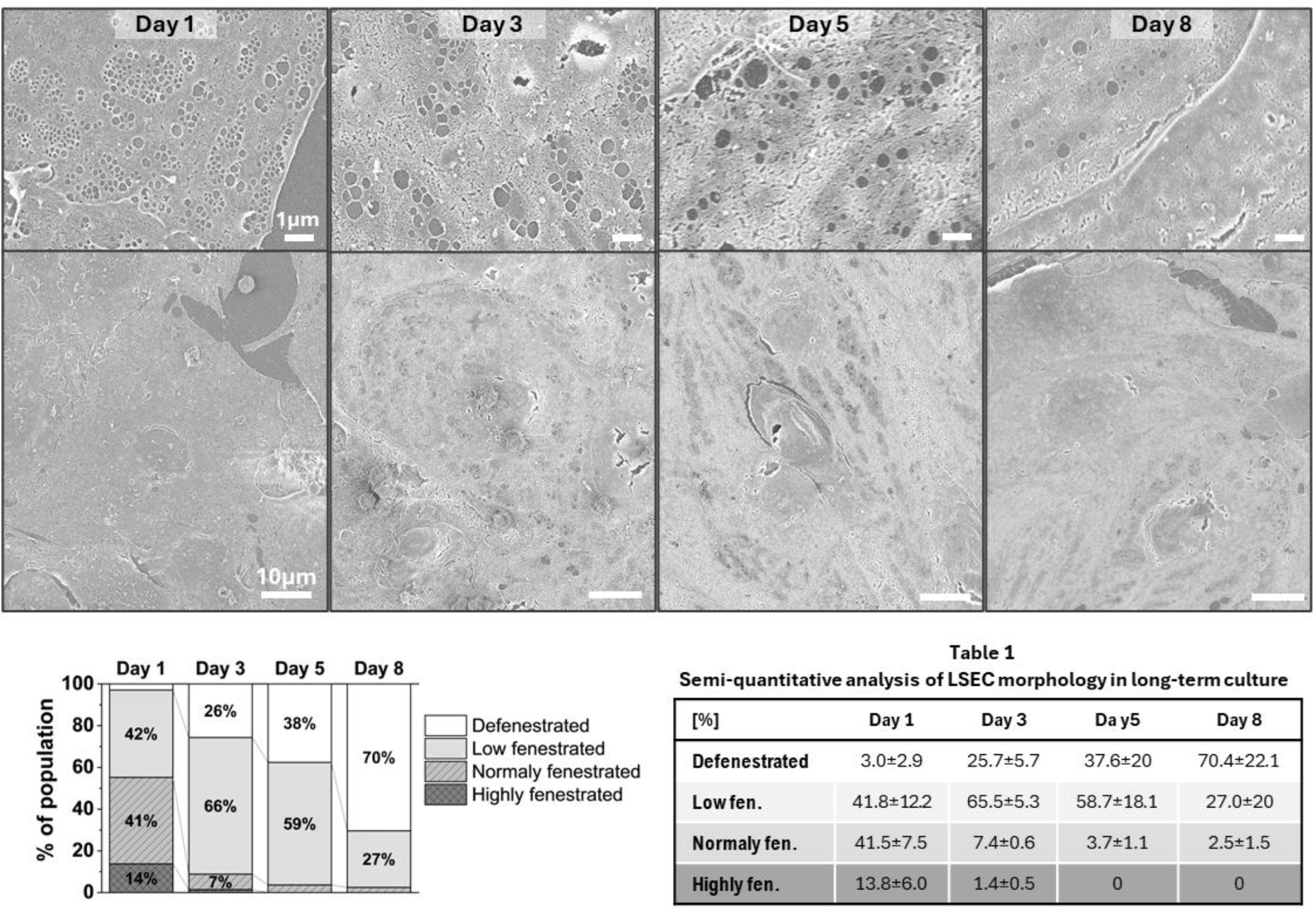
Ultrastructure of primary mouse LSEC over time in culture. Representative SEM images of LSEC cultured for 1, 3, 5 and 8 days. The top row presents higher magnification images of fenestrations showing gradual loss of organisation in the form of sieve plates and an increasing size of fenestrations over time. Semiquantitative analysis of the whole population confirmed the loss of fenestrations and increasing defenestration (complete loss of fenestrations) over time (n=6).

### 3.4. Changes in morphology and cytoskeleton of LSEC in long-term culture

Freshly isolated mouse LSEC display a characteristic cobblestone morphology with well spread monolayer of cells in nearly 100% confluency. The initial rounded shape of the cells becomes more elongated in the first days of culture (**Figure 4A**). LSEC maintain the characteristic for endothelial cell monolayers by compensating the decreasing number of the cells in the first 8 days of culture (**Figure 3A**) with increased cell size. The same morphological changes were observed throughout the whole sample suggesting a homogenous cell population. The elongation of the cell shape corresponds with changes in both actin and tubulin cytoskeleton which reorganized into characteristic parallel fibres supporting the elongated cell shape (**Figure 4B**). To evaluate how these cytoskeleton changes affect the fenestrated ultrastructure of LSEC, we used scanning electron microscopy (**Figure 5**).

### 3.5. Defenestration rate of LSEC in long-term culture

The semiquantitative analysis of LSEC ultrastructure was applied to reveal the fenestration status in the whole LSEC population and avoid cell selection bias. The purity of the cultures was confirmed by detailed analysis of LSEC ultrastructure, revealing that >95% of the cells on day 1 present fenestrated morphology (**Figure 5**). Furthermore, SEM imaging revealed that cells lose fenestrated morphology rapidly in the first 48h of culture, with only about 10% of the population maintaining normal (physiological) levels of fenestrations. Nevertheless, complete defenestration is occurring more slowly, with up to 50% of cells displaying some fenestrations even after 8 days in culture (**Figure 5**). The decrease in the number of fenestrations per cell is accompanied by a visible increase in the fenestration diameter. Until about day 5, the remaining fenestrations are mainly organised in the form of sieve plates (groups of fenestrations), while afterwards we observed primarily evenly distributed enlarged fenestrations.

## 4. Discussion

In this study, we performed detailed validation of an optimised long-term 2D monoculture system for primary mouse LSEC (e.g., high purity, monolayer confluency, 5% O_2_, EGM with half media volume exchanged daily) showing the preservation of cell viability and high endocytic capacity for at least 10 days. While the loss of fenestrations, a characteristic for LSEC dedifferentiation, still occurred as anticipated, the maintenance of the fenestrated phenotype was extended compared to other reports on 2D LSEC cultures [35], [36]. Moreover, the previously challenging preservation of high-throughput scavenging capacity allowed for the first successful application of *in vitro* siRNA-mediated knockdown in mouse LSEC monocultures using lipofectamine transfection.

Our results highlight a critical methodological consideration often overlooked in liver endothelial research: the rapid phenotypic drift of primary LSEC. The optimisation of long-term culture of primary LSEC has been ongoing for years in parallel with the attempts to create an LSEC cell line [22], [36], [37] . Stem cell derived LSEC are a promising alternative to primary cells, but preservation of fenestrated morphology in LSEC derived from human pluripotent stem cells (hPSC) still requires a maturation step by implantation into mice livers to expose hPSC-LSEC to the hepatic microenvironment [38]. Overall, all these attempts were met with some success, but still precluding the use of methods that require several days to develop or *in vitro* studies of prolonged exposure, such as hepatotoxicity in drug development. Nonetheless, some studies use primary LSEC cultured beyond the dedifferentiation time without validating their liver-specific identity. DeLeve and Maretti-Mira have notably argued that the lack of rigorous validation, specifically the demonstration of fenestrae via electron microscopy, constitutes a “breach of transparency” that undermines reproducibility in the field [21]. By coupling quantitative SEM analysis with functional assays, we demonstrate that while fenestrations are indeed lost within a few days due to the absence of the hepatic microenvironment, the endocytic function can be maintained. This distinction redefines the valid window for *in vitro* experimentation, suggesting that while >5-day-old cultures are unsuitable for studying porosity, they can remain a valid model for receptor-mediated endocytosis and intracellular trafficking studies if cultured in the right conditions.

The observed decoupling of fenestration maintenance and scavenger function in our model may be partially attributed to the mechanical environment. We cultured cells on fibronectin-coated plastic (with stiffness in the GPa range), which is orders of magnitude stiffer than the physiological liver sinusoid (~600 Pa - 1 kPa)[39]. Recent mechanobiology studies indicate that substrate stiffness is a primary driver of LSEC dedifferentiation: cells on soft hydrogels or in 3D spheroids retain fenestrae significantly longer than those on stiff glass or plastic [40][41][25]. While softer substrates might preserve fenestrated morphology better, they often restrict the number of methods that can be used to study LSEC in those systems. Thus, our optimised 2D LSEC monoculture represents a pragmatic compromise that enables robust imaging strategies with electron and fluorescent microscopy as well as high-efficiency transfection and quantitative assessment of scavenging functions.

We optimised the mouse LSEC long-term culture conditions based on the existing reports as well as years of in-house experience. The immunomagnetic separation of CD146-positive cells was proven to produce high yields of LSEC with purity >95% [28], [42]. High purity is crucial for long-term culture as any proliferating cell can contaminate and outcompete the endothelial cells. The seeding density and oxygen tension were also previously shown to be important for maintenance of LSEC phenotype and function, mimicking the physiological low-oxygen conditions of the liver sinusoids. Martinez et al. showed that low oxygen conditions (5% O_2_) significantly slow LSEC dedifferentiation and reduce oxidative stress [18]. In our study, we observed an increase in the reduction potential of the cell in the first 3-5 days of culture, which suggests that a decrease in oxidative stress may be crucial for LSEC *in vitro*. The culturing media and their composition also greatly influence the maintenance of LSEC phenotype and function. For example, the use of corticosteroids such as dexamethasone was shown to delay defenestration but did not affect the decline of scavenger function [43]. This study implemented full endothelial growth media widely used for endothelial with some reports suggesting a beneficial effect also in LSEC culture [35].

Until now, siRNA knockdown has been one of the techniques excluded from studying LSEC *in vitro* due to the rapid dedifferentiation of these cells. There are multiple *in vivo* studies demonstrating silencing of selected genes in animals injected either with siRNA directly [44] or with siRNA-containing nanoparticles [45]. As the successful transfection can take up to several days, only a few recent reports showed successful use of siRNA [46][47], all performed on human LSEC which are known to dedifferentiate at a slower rate than rodent LSEC [26], [48]. Our findings show that efficient gene silencing can also be achieved in pure mouse LSEC monocultures, avoiding the complexity and limitations of co-culture systems. The prolonged time in culture during which LSEC preserve their characteristics allowed us to demonstrate the Mrc1 *in vitro* knockdown also in mouse LSEC. This result opens new possibilities for the application of the method to detailed studies of molecular mechanisms of still unknown regulators of fenestrations [2], [48] and the scavenging system [3].

Interestingly, our siRNA knockdown experiments revealed an apparent functional discrepancy: while Mrc1 protein levels and RibB binding capacity dropped by ~70%, the degradation capacity only decreased by ~30%. A delay between RNA silencing and full loss of receptor protein is expected, given the relatively slow turnover of MRC1 [34]. A study in mannose receptor knockout mice (C57BL/6 background) showed that LSEC partly relies on mannose receptor-mediated endocytosis of lysosomal enzymes to keep up their high catabolism of endocytosed material [33]. These enzymes are released into blood from tissue turnover processes, and the enzymes’ function can be maintained for days after internalization in the LSECs [49]. The more modest decrease that we observed in the LSEC degradation efficiency of RibB than in the internalisation of the ligand can be explained by the high stability and long half-life of the lysosomal enzyme pool acquired by the cells prior to the knockdown. This lysosomal inertia suggests that for short-term experiments (<72h), LSEC maintain high degradation capability even when receptor-mediated enzyme recruitment is compromised. This finding has important implications for testing drug delivery systems, suggesting that LSEC scavenging capacity is highly resilient and potentially difficult to fully control.

## 5. Conclusions

This study addresses a critical need in the field for extended *in vitro* culture systems of primary LSEC that preserve functional phenotype beyond the typical 24 h window. Here we demonstrate that, in optimized conditions, the functional endocytic capacity can be maintained for at least 10 days, despite ongoing structural dedifferentiation. This extended functional viability provides a critical window for the application of siRNA-mediated gene silencing, a technique previously limited to complex co-culture systems or *in vivo* approaches. Efficient knockdown of Mrc1 at RNA and protein levels demonstrates the feasibility and general applicability of this platform for studying LSEC-specific molecular mechanisms. The observation that receptor-mediated endocytic function persists well beyond the loss of fenestrations defines a temporal dissociation that can be strategically exploited for mechanistic research. This work establishes a practical and accessible *in vitro* system for future studies of still-unknown regulators of LSEC fenestration architecture and signalling pathways involved in the scavenging system.

## Acknowledgements

The authors would like to thank Randi Olsen and Tom-Ivar Eilertsen from the Advanced Microscopy Core Facility at UiT for their electron microscopy expertise.

## 6.1. Financing

Open access funding for the article publication charges was provided by UiT The Arctic University of Norway (incl University Hospital of North Norway). This study was supported by the European Union’s Horizon Europe project: “DeLIVERy” EIC-2021-Pathfinder grant no. 101046928; “ImAgE-D” MSCA-DN grant no. 101119613 and the Research Council of Norway FRIPRO “PACA-Pill” grant no. 325446.

## 6.2. Authors contributions

Conceptualization K.Sz.,B.A., C.H., E.S., K.S., P.M; Methodology K.Sz., B.A., G.D., D.H., C.H., E.S., K.S., P.M.; Formal analysis E.S.; Investigation K.Sz., B.A., E.S., D.H., C.F., G.D.; Funding: P.M; Writing (original draft) K.Sz., B.A.; Writing (reviewing and editing) all authors.

## Notes

### Competing Interest Statement

The authors have declared no competing interest.

