## Supplementary information for "Long-term 2D monoculture of primary mouse LSEC preserves scavenging capacity and enables siRNA knockdown of Mrc1"

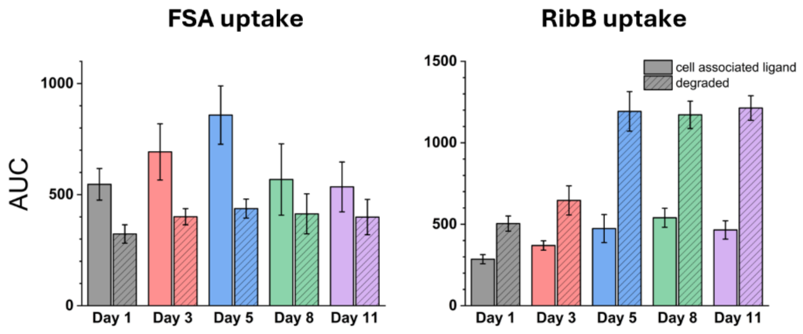


| dataset | Comparion between day X vs Y [ANOVA + Turkey, p-values] | | | | | | | | | |
|---|---|---|---|---|---|---|---|---|---|---|
| | 11 vs 1 | 11 vs 3 | 11 vs 5 | 11 vs 8 | 3 vs 1 | 5 vs 1 | 5 vs 3 | 8 vs 1 | 8 vs 3 | 8 vs 5 |
| FSA Cell ass. | 0.986 | 0.432 | 0.0109 | 0.934 | 0.495 | 0.0141 | 0.352 | 0.948 | 0.641 | 0.0235 |
| FSA degraded | 0.454 | 0.956 | 0.864 | 0.878 | 0.419 | 0.0991 | 0.906 | 0.261 | 0.918 | 0.894 |
| RibB Cell ass | 0.0012 | 0.152 | 0.892 | 0.375 | 0.258 | 0.0007 | 0.0989 | 9.89E-06 | 0.00204 | 0.465 |
| RibB degraded | 2.29E-10 | 2.42E-08 | 0.929 | 0.918 | 0.201 | 4.24E-10 | 4.84E-08 | 8.11E-10 | 8.96E-08 | 0.911 |

Figure S1. Area under curve (AUC) analysis of endocytic capacity data presented in Figure 1 and statistical analysis of comparison between AUC of endocytic capacity at selected days using one-way ANOVA with Tukey posthoc correction, significant values ($p<0.05$) are marked in green.

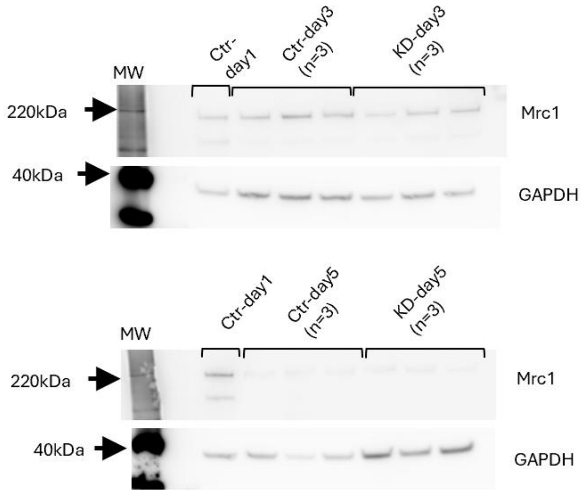


Figure S2 Raw images of the Western blot from representative experiments presented in Figure 2.
